# Evolutionarily conserved transcriptional landscape of the heart defining the chamber specific physiology

**DOI:** 10.1101/2021.01.22.427752

**Authors:** Shrey Gandhi, Anika Witten, Federica deMajo, Martijn Gilbers, Jos Maessen, Ulrich Schotten, Leon J. de Windt, Monika Stoll

## Abstract

Cardiovascular disease (CVD) remains the leading cause of death worldwide. A deeper characterization of the regional transcription patterns within different heart chambers may aid to improve our understanding of the molecular mechanisms involved in the function of the heart as well as our ability to develop novel therapeutic strategies. Here, we determined differentially expressed protein coding, long non-coding (lncRNA) and circular RNA (CircRNA) genes within various heart chambers across seven vertebrate species. We identified chamber specific genes, lncRNAs and pathways that are evolutionarily conserved in vertebrates. Further, we identified lncRNA homologs based on sequence, secondary structure, synteny and expressional conservation. Interestingly, most lncRNAs were found to be syntenically conserved. Various factors affect the co-expression patterns of transcripts including (i) genomic overlap, (ii) strandedness and (iii) transcript biotype. We also provide a catalogue of CircRNAs which are abundantly expressed across vertebrate hearts. Finally, we established a repository called EvoACTG (http://evoactg.uni-muenster.de/), which provides information about the conserved expression patterns for both PC genes and non-coding RNAs (ncRNAs) in the various heart chambers, and may serve as a community resource for investigators interested in the (patho)-physiology of CVD. We believe that this study will inform researchers working in the field of cardiovascular biology to explore the conserved yet intertwined nature of both coding and non-coding cardiac transcriptome across various popular model organisms in CVD research.

## Introduction

The vertebrate heart is a complex organ and has undergone remarkable anatomical changes during evolution. The anatomy of the heart varies drastically among vertebrates, with a two-chambered heart in fish, three-chambers in amphibians, and a complex four-chambered heart in mammals (Stephenson et al. 2017; Jensen et al. 2013). Even within organisms, the chambers of the heart follow different developmental trajectories and play distinct roles in maintaining cardiac function and homeostasis (Boogerd et al. 2009; Lin et al. 2012). Apart from the anatomical differences, cardiac regenerative capability also varies among vertebrates (Vivien et al. 2016). In recent years, several studies have explored the cardiac transcriptome across various organisms (Cardoso-Moreira et al. 2019; Sarropoulos et al. 2019; Necsulea et al. 2014), yet most of these studies ignored the local differences within the heart. Considerable heterogeneity exists within individual hearts, with each chamber demonstrating profound expression differences (Singh et al. 2016; Johnson et al. 2018). With the advent of next generation sequencing (NGS), it is now possible to identify the conserved molecular mechanisms underlying the development of vertebrate hearts.

In particular, long non-coding RNA (lncRNA) molecules have been shown to play a prominent role in cardiac development, expression regulation and pathophysiology of cardiovascular diseases (Gomes et al. 2017). LncRNAs are known to regulate various processes including epigenetic modifications, transcription, splicing, translation and the expression of miRNA and transcription factors (Mallory and Shkumatava 2015; Ma et al. 2015). However, little is known about the expression and conservation of these transcripts within the different compartments of the heart (Gandhi et al. 2019). Unlike protein coding (PC) genes, lncRNAs do not have conserved sequence similarity and rapid evolutionary turnover renders the identification of orthologs challenging. The few studies that investigated the conservation of lncRNAs found little evidence of sequence conservation (Necsulea et al. 2014), while other studies attempted to identify lncRNA homologs based on syntenic organization (Hezroni et al. 2015; Chen et al. 2016; Bryzghalov et al. 2020).

Here, we provide insights into the key pathways and expression modules that contribute to the conserved yet locally distinct nature of cardiac tissue within vertebrates in an attempt to gain a better understanding of the molecular architecture of the heart in health, adaptation and disease. We used bulk RNA-Seq to explore the expressional landscape of cardiac transcriptomes for seven vertebrate species representing different stages of the evolutionary development of the heart. We elucidated the regional diversity existing across heart chambers and detected novel myocardial lncRNAs and circular RNAs (circRNAs) for all organisms. Moreover, we determined homologous lncRNAs based on sequence, structure, and syntenic conservation. Additionally, we investigated which important factors influence the co-expression of neighboring gene pairs and, further, if the conservation of these pairs is important for overall gene regulation. Finally, we established a repository called EvoACTG (http://evoactg.uni-muenster.de/), which provides information about the conserved expression patterns for both PC genes and non-coding RNAs (ncRNAs) and may serve as a community resource for investigators interested in the (patho)physiology of CVD.

## Results

### Mapping the cardiac transcriptome

We generated gene expression data using strand-specific total RNA-Seq for seven vertebrate species namely zebrafish, African frog, chicken, rabbit, mouse, human and goat (Supplemental Table S1). On average of 47.9 million reads (standard deviation ± 9.47 million reads) was reached per sequencing library, with the mapping percentage varying between species likely reflecting the quality of the reference genomes (Supplemental Table S2). The average percentage of uniquely mapped reads varied between 89.9% in humans to 52.5% in goat samples. The goat samples also had the highest percentage of multi mapped reads with an average of 44.4% reads mapping to more than one locus.

### Identification and classification of lncRNAs

In each species, we identified novel spliced transcripts, which were subsequently evaluated for their protein-coding prowess to discover multi-exonic lncRNAs. On average 1,350 novel lncRNAs per species was detected, with a maximum of 3,366 lncRNAs in African frogs. Due to the different quality of genome assembly and asymmetric gene annotations there are large differences in the number of known lncRNAs across species (Supplemental Fig. S1). While there are no annotated lncRNAs in the rabbit genome in Ensembl, the human and mouse genomes had more than 45,000 annotated lncRNAs.

Next, we classified lncRNAs based on their genomic position relative to the nearest PC gene (Fig. 1). The percentage of intergenic lncRNAs ranged from 52 to 85 percent for species with a smaller number of annotated genes (chicken, frog, goat and rabbit). Notably, it was much lower for human (22.4%), mouse (19.9%) and zebrafish (39.8%) with well annotated genomes. The mouse, human and zebrafish genomes had the highest proportion of exonic lncRNAs, particularly nested sense lncRNAs. The percentage of intronic lncRNAs ranged between 5.1% in mouse to 22.6% in frogs. The differences in lncRNA categories among species may have some biological cause, but appears to be driven by the quality of the reference genome annotations.

**Figure 1:**
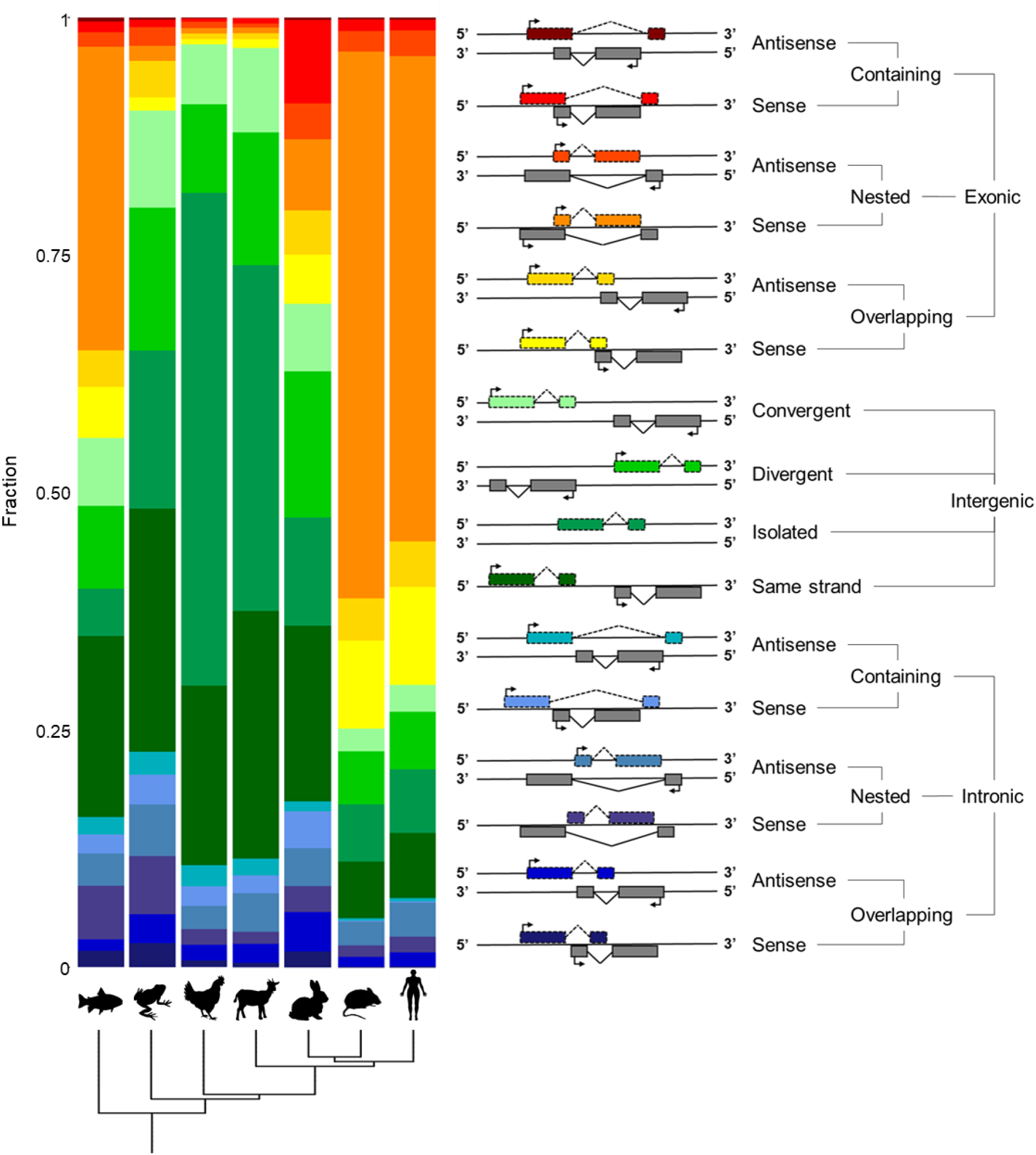
Distribution of various classes of lncRNAs. The graph depicts the relative distribution of various classes of lncRNAs. The lncRNAs were divided into various categories based on overlap with the closest coding gene. The right side of the panel represents the different classes (color coded) and their distribution is shown on the left side.

### Expression signatures differ across various heart chambers

We profiled the cardiac transcriptome of all organisms to identify myocardial genes and transcripts (Fig. 2A). The number of expressed genes ranged from 18,392 in frog to 11,581 in rabbit samples (Fig. 2B). Overall, the percentage of expressed genes varied between 45 to 48 percent for all organisms except human and mouse, where it was around 24 percent due to the higher number of annotated genes. Next, we examined the distribution of expressed genes according to their biotype. While the percentage of expressed PC genes was between 48 to 58 percent, the percentage fluctuated considerably for lncRNA genes owing to the annotation disparity among species.

**Figure 2:**
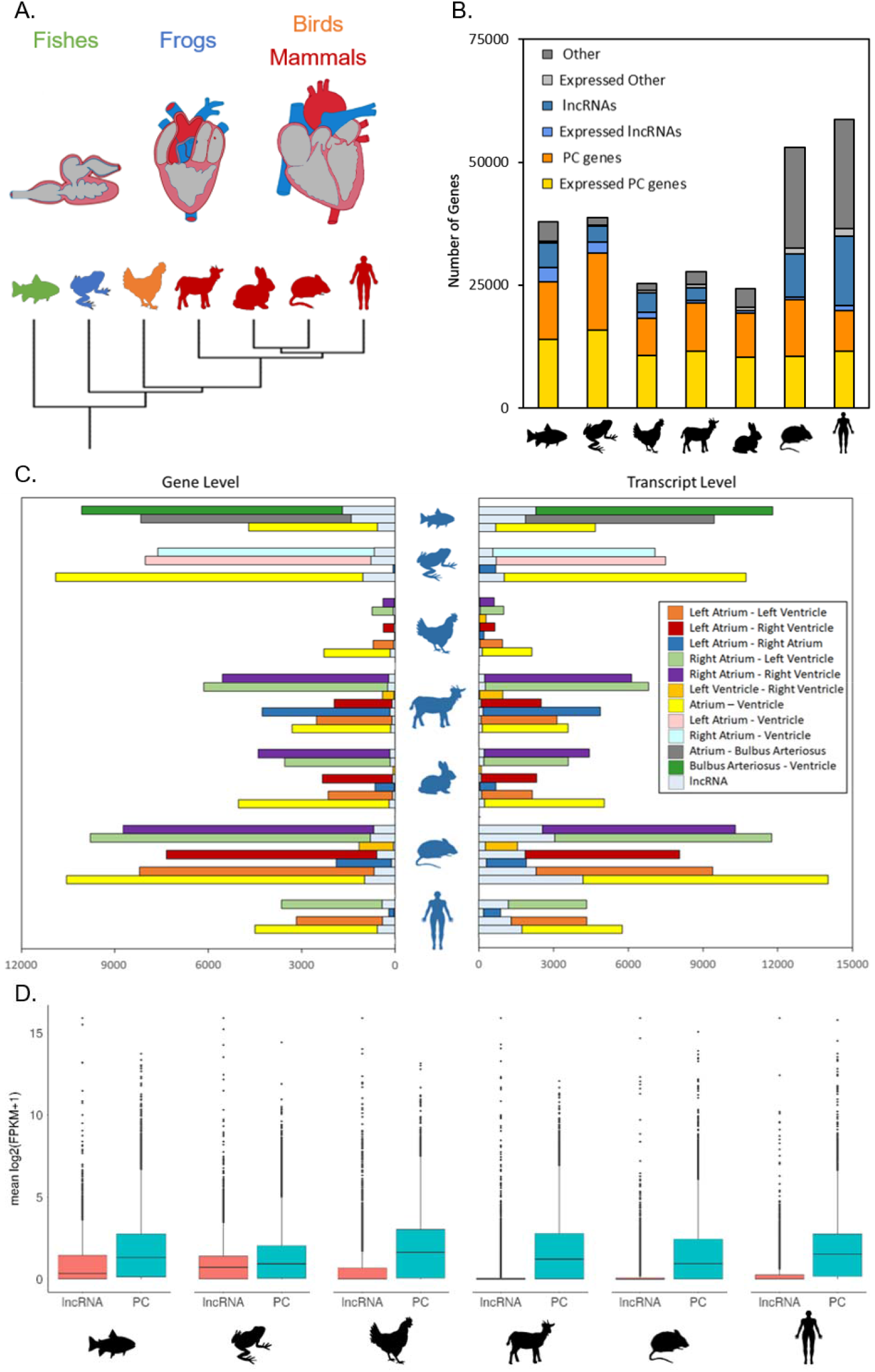
Expression signatures differ across various heart chambers. (A) Cartoon representation of heart morphologies across vertebrates. (B) Distribution of PC, lncRNA and other cardiac expressed genes. (C) Number of DEGs and DETs across each species. (D) Average gene expression level of lncRNA and PC genes.

Next, we performed principal component analysis (PCA) using variance-stabilizing transformation (VST) read counts for the top 500 genes based on the chamber biopsy details. In all species, PCAs were distinctly able to separate the atrial and ventricular samples (Supplemental Fig. S2). The zebrafish heart samples including bulbus arteriosus, clearly grouped according to the tissue biotype. For goat, mouse and rabbit samples, the differentiation between left (LA) and right atrial (RA) biopsies was more evident than between the ventricles. In chicken and frog, the samples appear to group by organism rather than LA and RA. Also, in humans, the samples seemingly grouped by individuals, but an effect of gender was also apparent. We reanalyzed the human chamber specific data based on the study by Johnson et al. and observed similar segregation of samples based on the gender (Johnson et al. 2018).

To examine the regional differences in expression patterns, we individually performed differential expression analysis for all possible heart chamber comparisons across each species. Only genes/transcripts with an absolute fold change ≥ 1 (adj. p-value < 0.05) were considered as differentially expressed genes (DEGs) /transcripts (DETs). The least number of DEGs were observed in chicken, possibly limited by the sample size. For all species, the number of DEGs within ventricles and atria were larger than the differences between them (Fig. 2C). Except for chicken, for all organisms with four chambered hearts, the number of DEGs was slightly higher when comparing the right half of the heart than the left part, in line with previous studies (Johnson et al. 2018). Similarly, there were more DEGs expressed between the two atria than the two ventricles, with the exception for chicken. The number of differentially expressed lncRNA genes and transcripts varied significantly among all species. Although we detected several conserved DEGs across the cardiac chambers for all vertebrates, there were also many DEGs that were unique to comparisons within each species (Supplemental Table S3A-G).

Next, we compared average gene expression levels for lncRNA and PC genes across species. The expression levels were significantly higher for PC genes than lncRNAs for all species considered in the analysis (Wilcoxon signed-rank test, p-value< 2.2e-16). (Fig. 2D). We did not consider rabbit samples for this analysis, since we only have novel lncRNA annotations, which were determined using fragments per kilobase of transcript per million mapped reads (FPKM) value > 0.5 in at least 2 samples.

### Several genes express in a chamber-specific manner

For each species, we determined the tissue specificity using Tau (τ) values of all the expressed genes within our dataset (Yanai et al. 2005; Kryuchkova-Mostacci and Robinson-Rechavi 2017). Genes with τ score >= 0.65 for any of the heart chambers were considered to be expressed tissue specific. While most of the myocardial genes were expressed ubiquitously in the heart, genes such as *NPPA, MYBPHL* and *MYL2*, which are known to be chamber specific, also demonstrated tissue specificity in our dataset (Supplemental Fig. S3A) (Asp et al. 2012; Song et al. 2015; Barefield et al. 2017). In accordance with the differential expression results, most of these chamber specific genes were expressed in the atria. Within the atria, the majority of these genes were expressed in RA. The number of chamber specific PC and lncRNA genes were equally proportional to the number of cardiac expressed genes (Supplemental Fig. S3B). Notably, several genes including *PITX2, IRX4* and *BMP10* exhibited conserved chamber specific expression patterns for LA, ventricle and RA, respectively, across all the seven species.

Next, we looked at the expression of known cardiac transcription factors (TFs) and checked if their expression was conserved across species (Supplemental Table S4). For most of the cardiac TFs including *HAND2, GATA6* and *NKX2-5*, we observed broad expression profiles across the heart in all the organisms. While *IRX4* and *TBX5* were expressed in a tissue specific manner in the ventricles and atria respectively across all the seven vertebrates. Several immune genes have been shown to be enriched in human atria (Johnson et al. 2018). We observed that, many immune genes including *CCL2, CCL3, CCL4, CCL8, CXCL1, CXCL2, CXCL3, CXCL4, CXCL8, CXCL14*, IL14, *IL16, IL18, IL15, IL10* and *IL4R* were found to be enriched in human atria, but their expression was not conserved across other vertebrates. Genes such as *CXCL16, IL1B, IL6* and *IL1R1* were expressed specifically in the atria across multiple vertebrates.

We also looked at the regional differences in the expression of ion channel genes and found most of these genes to be ubiquitously expressed in the heart. While the ion channel subunit genes *including KCNH6, KCNH7, KCNJ3, KCNJ5, KCNK1, KCNK3* and *CACNA1D* were enriched in the atria, others such as *KCNJ2, KCNJ8, CACNA2D1, SCN3B*, and *SCN4B* were specific for ventricles across most of the species. We also observed many genes, including *CACNA2D2, KCNA4, KCNA5, SCN1B*, and *SCN4B* whose tissue specificity varied across vertebrate species. There were several ion channel genes, including *HCN4* and *HEY2*, which demonstrated chamber specific expression across vertebrates (Supplemental Table S5). These results demonstrate regional expressional differences existing within vertebrate hearts. The fact that these differences are conserved indicates the preserved nature of larger biological programs functional within the heart.

### Expression modules enriched in various cardiac chambers are conserved

To determine the pathways and expression modules enriched in the different heart chambers, we performed gene set enrichment analysis (GSEA). Using DEGs between various chambers, we identified several significantly enriched gene ontology (GO) terms. These enriched terms were grouped in enrichment clusters which revealed different biological processes, molecular functions and cellular components active in the heart chambers. Concordant with differential expression, enrichment analysis revealed multiple conserved enriched GO terms when comparing LA and RA and the ventricles. In most organisms, comparison between LA and RA revealed multiple clusters of terms involved in metabolic processes, energy metabolism, and heart valve development (Supplemental Fig. S4A). Cellular component clusters mostly corresponded to mitochondria and ribosomes. Similarly, for left ventricles (LV) and right ventricles (RV), we found enrichment of terms mapping to cardiac muscle contraction and respiratory chain complexes (Supplemental Fig. S4B). We found no enrichment for mitochondrial complexes in the ventricular comparison.

Next, we examined the differences between the left and right side of the heart. For LA and LV, most GO terms clusters were similar to the ones obtained in the earlier comparisons, with the exception of fatty acid metabolism, tricarboxylic acid cycle and peptidase activity (Supplemental Fig. S4C). Furthermore, the comparison of the right side of the heart revealed enrichment for terms involved in BMP signaling and heart valve development (Supplemental Fig. S4D). Also, these enrichment clusters were more conserved for the comparison between the heart sides than for LA vs RA and LV vs RV.

Next, we looked at the enrichment of terms within each chamber against the average expression of all the other chambers for this species. Through this, we were able to identify terms that were either exclusive or universally enriched across each chamber. In all species, except zebrafish, we found most enriched terms in the RA compared to other chambers. In zebrafish, the bulbus arteriosus displayed the maximum number of enriched terms. We detected several conserved clusters involved in energy metabolism, respiratory processes, mitochondrial and ribosomal assembly that were enriched across all chambers, but the enrichment profiles differed profoundly. Most of the terms involved in these clusters were negatively enriched in both the atrium, while for the ventricles they were positively enriched.

In the case of LA, we were able to detect several unique terms that were positively enriched for chemokine binding and leukocyte chemotaxis. (Supplemental Fig. S5A). Most of the positively enriched terms unique to LV were related to peptidase activity, while BMP signaling and heart development terms were negatively enriched (Supplemental Fig. S5B). The RA enrichment profile consisted of terms positively enriched in heart development, ion channel activity, WNT and BMP signaling pathways (Supplemental Fig. S5C). Finally, most of the terms were positively enriched for RV, including the processes involving fatty acid metabolism and methyl transferase activity (Supplemental Fig. S5D). These enrichment results illustrate the conserved nature of the spatial differences existing within vertebrate hearts.

### Sequence based conservation in lncRNAs is minimal

LncRNAs are not well conserved at the sequence level (Necsulea et al. 2014; Hezroni et al. 2015), yet they share short stretches of sequence conservation, which can be essential for their proper function. We only investigated the conservation of intronic and intergenic lncRNAs to remove the conservation bias, which may exist due to PC genes. Using blastn and OrthoMCL, we built homologous lncRNA families and found very few lncRNAs with sequence conservation across all species (Supplemental Table S6A). Most of the lncRNAs were found to be paralogs and not across different species. We applied the same strategy on the promoter regions of lncRNAs since previous studies reported the promoter regions of lncRNAs to be more conserved (Carninci et al. 2005). Our data support the previous reports indicating a poor conservation of lncRNAs at the sequence level. (Supplemental Table S6B).

### Most lncRNAs are conserved by syntenic location

Recent studies have shown that several lncRNAs have conserved syntenic location (Herrera-úbeda et al. 2019; Bryzghalov et al. 2020; Amaral et al. 2018). Hezroni et al. were also able to detect several syntenically conserved lncRNAs even when the sequence was not conserved (Hezroni et al. 2015). Latos et al. demonstrated that the sequence and the length of lncRNA *Airn* was inconsequential and only the positional transcription from the overlapping region with the *Igf2r* gene was important for its function (Latos et al. 2012). Therefore, we searched for lncRNAs whose position was conserved in relation to neighboring PC genes. We considered lncRNAs for which either the entire surrounding locus or the immediate PC genes could be consequential for expressional regulation. Based on the conservation of the orientation of neighboring genes we then classified lncRNAs into stranded or unstranded syntenic homologs (Supplemental Table S7A-B).

We detected 11,480 human lncRNAs with 7,386 lncRNAs unstranded syntenic homologs in the mouse. We also identified 71 human lncRNAs, which had one or more syntenic homologs in all the other 6 species, none of which were stranded syntenic homologs. For immediate syntenic homologs, the majority of the neighboring PC genes had the same orientation in both species, unlike the ones for which we checked larger background loci.

Next, we checked whether these syntenic homologs have more sequence conservation compared to non-syntenic random lncRNAs. For this, we considered the sequence conservation of human-mouse syntenic pairs and compared it with random human-mouse non-syntenic lncRNA conservation. We found that across all categories, syntenically conserved human-mouse lncRNAs had significantly more sequence conservation (Wilcoxon signed-rank test, p-value=4.08e-47). Also, there was little difference between the sequence conservation between stranded and non-stranded syntenic pairs (Supplemental Fig. S6). These results demonstrate the strong positional conservation of lncRNAs across species. This conservation might be a result of the conserved functional association of these lncRNAs with their neighboring lncRNA counterparts (Engreitz et al. 2016).

### lncRNAs with conserved secondary structure are rare

Selection pressure may be acting both at the sequence and structure level (Pegueroles and Gabaldón 2016). Despite low sequence conservation, few lncRNAs such as *HOTAIR, NEAT1* and *XIST* have been demonstrated to have a conserved secondary structure, which is deemed important for their function (Somarowthu et al. 2015; Lin et al. 2018; Pintacuda et al. 2017). Therefore, we attempted to identify lncRNA homologs based on secondary sequence conservation. We only considered lncRNAs shorter than 1,000 bases, since with increasing length prediction accuracy becomes less reliable. We discovered numerous lncRNAs with short regions of structural conservation across species, when focusing on lncRNA homologs with a structural identity >50% (Supplemental Table S8). We also identified several lncRNAs with a conserved secondary structure even when the sequence was not as conserved. For humans and mice, we found 101 conserved lncRNAs with structural identity greater than 50 percent. Of these, only 56 human-mouse homologs had a sequence identity greater than 40 percent. Although we do not observe strong structural conservation across lncRNAs, this may be driven by the accuracy of the structural prediction and alignment tools.

### Several factors influence the co-expression patterns of lncRNA and PC genes

LncRNAs are known to act as cis-regulators of gene expression and modulate the expression of neighboring genes. To investigate the effect of lncRNA expression on adjacent genes, we focused on robust human and mouse genomes and calculated the correlation coefficients for all transcript pairs within a 100 kb radius. Since previous studies have shown that the distance between genes influences the expression correlation values (Sarropoulos et al. 2019), we were interested in examining the factors which influence the co-expression value. We observed that transcript pairs having overlapping genomic positions had a significantly higher correlation coefficient than non-overlapping ones in both human (Wilcoxon signed-rank test, p-value= 1.08e-59) and mouse (Wilcoxon signed-rank test, p-value= 2.42e-117) samples (Fig. 3A). There was no significant difference between gene transcripts with partial overlaps and completely imprinted transcript pairs.

**Figure 3:**
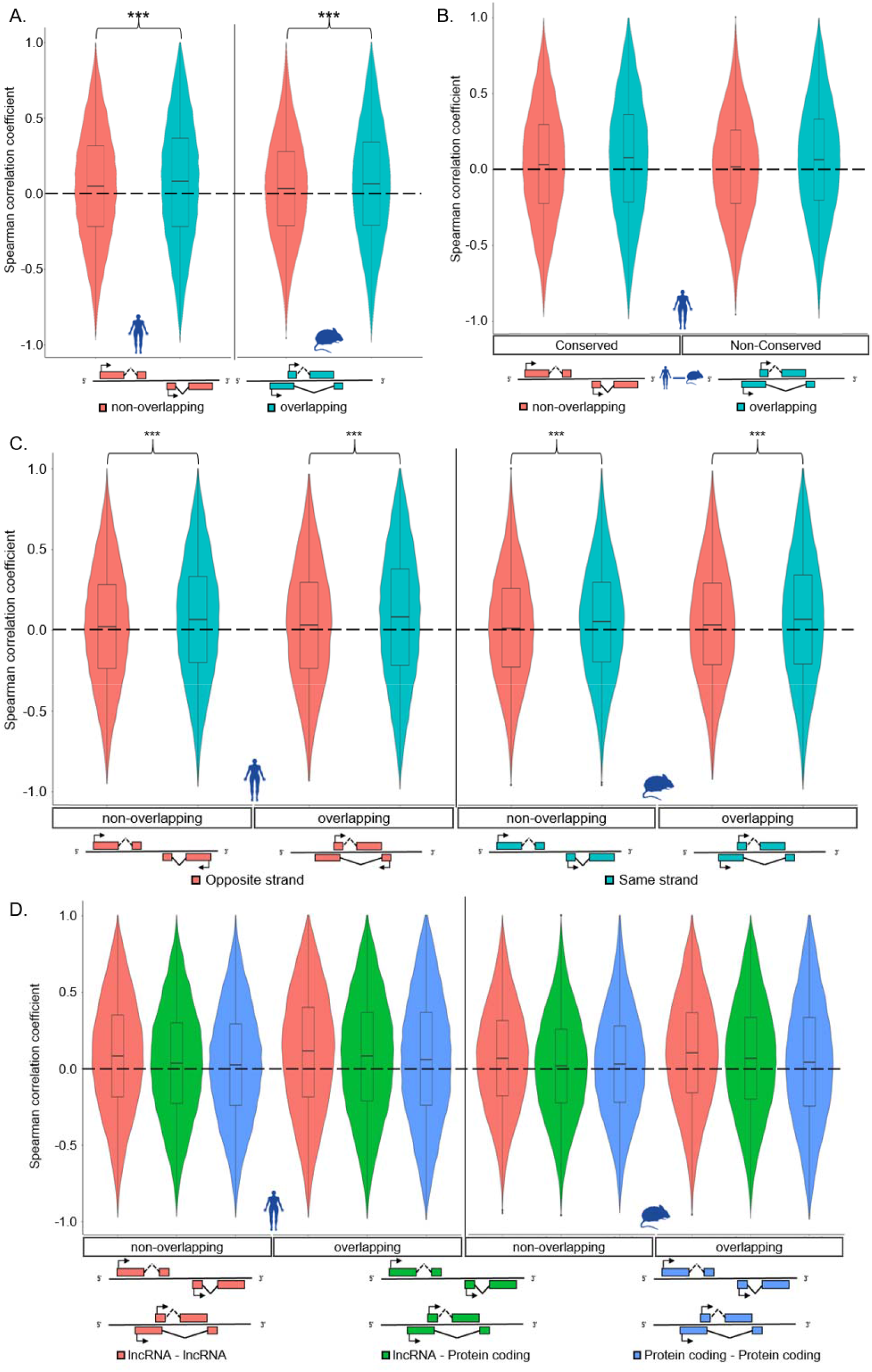
Several factors influence the co-expression patterns of lncRNA and PC genes. The violin plot shows the distribution of Spearman correlation coefficient for (A) overlapping and non-overlapping transcripts. The overlapping transcripts have significantly higher correlation than non-overlapping for both human (p-value=1.08e-59) and mouse (p-value=2.42e-117). (C) The transcripts on the same strand are more positively correlated than the ones on opposite strand both for human and mouse; ***= (Wilcoxon signed-rank test, p-value< 2.2e-15). (D) The correlation coefficients for lncRNA-lncRNA, lncRNA-PC and PC-PC pairs were significantly different for both overlapping and non-overlapping transcripts. The statistical analysis was done using Kruskal-Wallis test followed by Dunn’s test to analyze the multiple group comparisons. (B) The distribution of correlation coefficient for conserved mRNA and syntenically conserved lncRNA pairs between humans and mouse. The distribution has been plotted for human samples which show significantly higher correlation coefficients for the conserved transcript pairs compared to non-conserved lncRNA-PC pairs, both for overlapping and non-overlapping transcripts.

Next, we considered the strandedness of the transcripts and whether this has an effect on co-expression values. We observed that irrespective of the type of genomic overlap, transcript pairs on the same strand have significantly higher positive correlation than the ones on the opposite strand for both human and mouse samples (Wilcoxon signed-rank test, p-value< 2.2e-15) (Fig. 3C). We also observed significant differences in correlation values for lncRNA-lncRNA, lncRNA-PC and PC-PC transcript pairs (Kruskal-Wallis test, p-value< 2.2e-45) both for overlapping and non-overlapping neighbors. We detected highly positive correlation values for lncRNA-lncRNA pairs, followed by lncRNA-PC and least positive for PC-PC for all pairs of transcripts in humans (Fig. 3D). The exception being in non-overlapping transcripts of mice, where the lncRNA-PC pairs had the least positive mean correlation coefficient (Supplemental Table S9).

We then investigated whether the correlation coefficient is different for evolutionarily conserved gene pairs between human and mouse samples. There was no significant difference in the distribution of the correlation coefficient between conserved and non-conserved PC-PC gene pairs. For lncRNA-PC pairs, we looked for conserved syntenic lncRNAs and their conserved PC gene partners. The mean correlation coefficient of conserved syntenic lncRNA-PC gene pairs was significantly higher than that of non-conserved ones both for overlapping (Wilcoxon signed-rank test, p-value= 0.0016) and non-overlapping transcripts (Wilcoxon signed-rank test, p-value= 9.32e-15) (Fig. 3B). These results demonstrate that neighboring genes, in particular the overlapping genes are more positively co-expressed. Additionally, we determine several factors including the genomic overlap, strandedness, transcript biotype and conserved nature of the transcripts, which influences the co-expression of neighboring transcript pairs.

### CircRNAs are abundantly expressed in the heart

Several studies have shown that circRNAs are abundantly expressed in the heart and play an important role in cardiovascular pathophysiology (Werfel et al. 2016; Garikipati et al. 2019; Tan et al. 2017). Recently, cardiac circRNAs have also been shown to code for micropeptides (van Heesch et al. 2019). We therefore aimed to identify novel circRNAs and study their expression profiles across heart chambers for all seven species. We detected hundreds of novel circRNAs expressed in all seven species (Fig. 4A). While most of the circRNAs were exonic, we also detected a small number of intronic and intergenic circRNAs (Fig. 4B). Some genes such as *RYR2, MLIP* and *CORIN* produced multiple circRNA isoforms in several species. While *MLIP* and *RYR2* produced multiple circRNA isoforms in mammals and chicken, we detected several *CORIN* circRNA isoforms across all organisms. The maximum number of circRNA isoforms in human and mouse samples, originated from the *TTN* gene with 16 isoforms in mouse and 59 isoforms in human samples, in accordance to previous observations that human orthologues in general produce more circularized transcripts (Aufiero et al. 2018).

**Figure 4:**
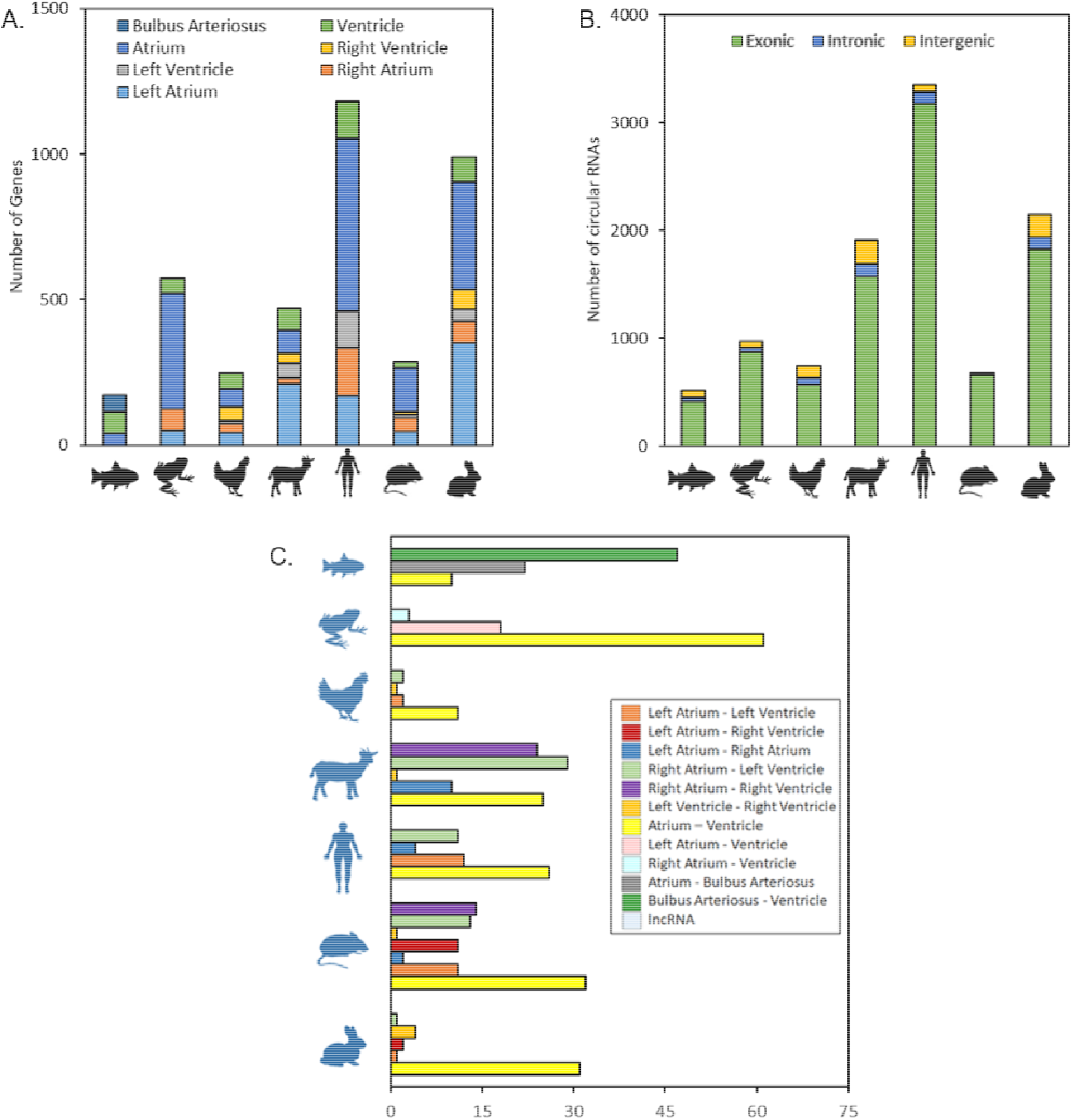
CircRNAs are abundantly expressed in the heart. (A) The distribution of chamber specific circRNAs across seven vertebrates. (B) The distribution of the biotypes of chamber specific circRNAs and (C) Number of differentially expressed circRNAs across each species.

Next, we examined the differential expression of circRNAs across heart chambers for all species (Supplemental Table S10). Similar to PC and lncRNA genes, the maximum number of circRNAs were differentially expressed between atria and the ventricles (Fig. 4C). For zebrafish, the bulbus arteriosus had the most differentially expressed circRNAs. We detected very few differentially expressed circRNAs when comparing within the atrial and ventricular biopsies.

We also identified circRNAs, in which the ratio of linear to circular isoforms for the same gene varied independent of each other across samples (Supplemental Table S11). One such circRNA originated from the RYR2 locus in human (1:237566567|237569319), mouse (13:11759671|11785141) and goat (28:35026494|35037789) samples, where the circular to linear transcript ratio differed between atria and ventricular biopsies. Another conserved circRNA originates from the *BNC2* gene in human (9:16435555|16437524) and goat (8:27805904|27807873) samples, where the circular to linear reads ratio is significantly higher in atrial samples.

We then calculated the tissue specificity of these circRNAs and detected several isoforms for genes such as *CORIN, RYR2*, and *RABGAP1L*, which exhibited tissue specific expression across species. The *CORIN* circRNA isoforms were specifically expressed in the atria in mammals, while in other species these isoforms were specific for ventricular tissue. *RYR2* circRNAs showed a higher tissue specificity score in ventricles for all species with a 4-chambered heart. *RABGAP1L, RERE, EVI5* circRNAs were exclusively expressed only in the atria of all mammals.

Next, we looked at the sequence conservation of circRNAs using a reciprocal blast hit (RBH) strategy and identified several homologs across species (Supplemental Table S12). Most of the homologous circRNAs were exonic circRNAs, with only 1 human circRNA, 15:98707562|98708107 from the *IGF1R* gene having homologs across all organisms. Several human genes such as *BNC2* and *RERE* had detectable circRNA homologs in all species with 4-chambered hearts.

Finally, we also identified pathways enriched in the genes that give rise to circRNA transcripts. Although most of the significantly enriched terms were only detected in humans, we were still able to detect a few enriched pathways in rabbit and mouse. The prominent enrichment term clusters include chromatin modification and organization, signaling pathways, biomolecular binding, signal transduction, regulation of biosynthetic processes and cardiac development among others (Supplemental Fig. S7).

## Discussion

Due to the complex nature of the heart as well as for ethical considerations, it is very difficult to obtain and analyze human biopsies. Most of the research in the cardiovascular field is hence performed in animal models, especially in mice whose cardiac anatomy differs considerably from humans (Wessels and Sedmera 2004; Lossi et al. 2016). Additionally, most of the transcriptome studies in the heart ignore the regional differences that exist within the heart. The lack of reproducibility of differential expression studies for CVDs has been attributed to several factors including the differing anatomy, differences in the genome quality, lack of uniform gene annotations and the absence of techniques to identify homologous transcripts, particularly for the non-coding genome. Here, we not only explore the temporal differences within the adult hearts of seven different vertebrate species, we also determine how conserved these differences are across these species. We describe the various expression modules active within each chamber of the heart and recognize conserved biological pathways exclusive to each heart chamber. We identify thousands of novel lncRNAs and explore the conservation of expressed lncRNAs based on sequence, secondary structure and syntenic position across these vertebrates. Additionally, we establish the various factors influencing the co-expression profiles of neighboring transcript pairs, including the genomic overlap, strandedness and the transcript biotype. Finally, we identify several novel circRNAs and explore their properties in the vertebrate hearts.

The complex nature of CVDs makes it difficult to study the underlying genetic mechanisms responsible for the disease. At the center of the cardiovascular system is the heart, which has undergone complex biological, anatomical and physiological changes across vertebrates and during the course of evolution. To understand these differences, we have to appreciate the conserved, yet dynamic diversity in the genetic pathways active inside the vertebrate hearts. Previous studies have focused on the chamber specificity of a few cardiac genes in the context of individual organisms (Lin et al. 2014; Kahr et al. 2011). These studies rarely consider all the heart chambers within the organisms, focusing only on the expressional differences for PC genes across particular heart chamber pairs. In this study, we calculate the chamber specific expression for all the genes, and considering all the heart chambers across seven vertebrate species. Using the expression data, we were able to show the conserved expression of several genes, including TFs like *IRX4* and *TBX5*, which are conserved across all species. We also identified several known chamber specific genes including *HCN4* (Garcia-Frigola et al. 2003), *BMP10* (Kahr et al. 2011), *PITX2* (Kirchhof et al. 2011), *TNFRSF12A* (Synnergren et al. 2020), *MYL2* (Asp et al. 2012) and *RPL3L* (Bond et al. 2019). We also determined several chamber-specific enriched terms conserved across all the vertebrates. While both the ventricles were enriched for mostly similar terms pertaining to respiratory processes, mitochondrial activity and energy metabolism, the biological processes enriched in the atria differed drastically. We found that LA was positively enriched for processes related to leukocyte chemotaxis, and RA was involved in heart development, WNT and BMP signaling pathways.

These results suggest an inherently conserved biological program, functional within each individual chamber of the vertebrate hearts. Genes such as *PITX2* are known to maintain left-right atrial identity, is known to be differentially expressed in the left atria of amphibians and mammals (Desgrange et al. 2018; Guerra et al. 2018; Franco et al. 2017). Our data indicates that *PITX2* is strongly enriched within the zebrafish atria, which is located on the left symmetry with respect to the ventricle, thus maintaining its lateral specificity. These conserved genes could also help determine the evolution of the heart morphology. In addition, there also exist regional differences within the hearts of individual organisms. Using differential expression analysis, we determined the regional differences within the individual heart chambers. These differences are partly due to the different origin and developmental trajectories each cardiac chamber undergoes.

While the dysregulation of PC genes is an important indicator of several CVDs, the focus has now shifted towards the ncRNAs. In the past decade, genome-wide association studies (GWAS) have identified a large number of loci mapping to the non-coding regions of the genome rather than pointing towards PC genes.

The identification of majority of the GWAS signals in the non-protein-coding region of the genome indicates a complex regulatory network driven by this unexplored epigenetic layer of gene regulation. The unavailability of well annotated ncRNA genomes, restricts the analysis to only known transcripts in some species. Here, we identify several novel lncRNAs, many of which are differentially expressed across the various chambers in each species. Despite the promise of functional genomics, the lack of evolutionary conservation and difficulty in identifying lncRNA homologs makes it difficult to translate the findings from animal models to humans.

To this end, we identify novel lncRNAs expressed within the various chambers of the heart for all the species we studied. Since most of the sequence based conservation methods have been designed for coding genome and lncRNAs are not that well conserved at the sequence level, our study also focused on the various other dimensions of evolutionary conservation. Rather than depending on sequence conservation, we established bioinformatics pipelines to identify lncRNA homologs based on the structure, synteny and expression conservation across the vertebrates. In the process, we also discovered lncRNA homologs based on these dimensions for the cardiac transcriptomes of seven vertebrates. Our study indicates that most of the lncRNAs are syntenically conserved across species even when the sequence or secondary structure is not. We also determine that several syntenically conserved lncRNAs demonstrate chamber specific expression. The conserved synteny and expression of these lncRNAs might be related to their conserved functional role pertaining to the neighboring PC genes.

Our investigation of the co-expression patterns of the neighboring genes demonstrated that these transcript pairs are significantly more correlated than random gene pairs. Our results also indicate that the overlapping genes had more correlated expression than the non-overlapping neighbors. Additionally, we found that the strandedness of these neighboring transcripts appears to influence the co-expression values, with genes on the same strand driving towards positive values. Our results suggest that the biotype of these neighboring transcript pairs also has an influence on the correlation coefficient with lncRNA pairs having maximum and PC pairs with minimum correlation coefficients. We also find that syntenically conserved PC-lncRNA pairs are highly correlated. These results indicate that not only the sequence but the genomic location of the transcripts, especially for the lncRNA genes, play an important role for their expression.

Additionally, we discovered that circRNAs are abundantly expressed across vertebrate hearts. We explored the conserved nature of the novel circRNAs across the cardiac tissue and identify several conserved circRNAs originating from important cardiac genes such as *MLIP* and *RYR2*. Most of the circRNA genes were found to be involved in regulatory processes such as chromatin modification and organization, signaling pathways, biomolecular binding, signal transduction and regulation of biosynthetic processes.

In conclusion, here we address the variability in the lncRNA annotations across popular animal models in CVD research. We tackle the problem of identification of lncRNA homologs by exploring the various dimensions of conservation and indeed find most lncRNAs to be syntenically conserved. We demonstrate several factors, which may influence the co-expression of transcripts including distance, overlap, strandedness and biotype. We hope that the exploration of the cardiac transcriptome of popular animal models will serve as a stepping stone to facilitate the investigation into the mechanistic role of various ncRNAs in CVDs. With this study and the corresponding database (http://evoactg.uni-muenster.de/), our aim is to enable researches to make an informed choice about the animal models to study based on the expression of the established coding and lncRNA genes involved in various CVDs.

## Methods

### Tissue acquisition, transcriptome sequencing and assembly

The chamber specific heart biopsies were obtained for seven vertebrate species including zebrafish (*Danio rerio*), African frog (*Xenopus laevis*), chicken (*Gallus gallus*), rabbit (*Oryctolagus cuniculus*), goat (*Capra hircus*), mouse (*Mus musculus*) and human (*Homo sapiens*). Total RNA was isolated from each of the sample biopsies, followed by the assessment of the quality of RNA. Directional bulk RNA Seq libraries were prepared after rRNA depletion and were then sequenced in paired-end mode 75 cycles on a NextSeq 500 system (Illumina). The resulting sequencing reads were trimmed using Trimmomatic (v0.36) (Bolger et al. 2014) followed by alignment to the respective reference genomes using the STAR aligner (v2.5.3a) in 2-pass mapping mode (Dobin et al. 2013). The mapped reads were assembled into transcripts using StringTie (v1.3.4d) guided by the reference genome annotations (Pertea et al. 2015) in the novel transcript identification mode.

### Identification and classification of cardiac expressed lncRNAs

Novel transcripts identified by StringTie were pre-filtered using different criteria, including length < 200 bases and FPKM > 0.5 in at least 2 samples, before using tools to identify and shortlist lncRNA transcripts. We employed CPC2 (v0.1) (Kang et al. 2017), CNCI (v2) (Sun et al. 2013), NCBI’s ORFfinder (v0.4.3) and Pfam database (RD et al. 2016) to screen the candidate transcripts. The resulting high-confidence novel lncRNAs were further classified based on the genomic location with respect to the PC genes using the FEELnc (v1.0) classifier (Wucher et al. 2017).

### Chamber specific expression of cardiac transcriptome

The novel lncRNA annotations were merged with the reference genome and were quantified again using StringTie. DESeq2 (v1.22.2) (Love et al. 2014) was used to identify DEGs/DETs for all possible chamber comparisons including combined atrial and ventricular contrasts for each species. Genes/transcripts with log2 fold change (FC) ≥ 1 or ≤ -1 and adjusted p-value < 0.05 (Benjamini-Hochberg) were classified as differentially expressed. The τ index (Kryuchkova-Mostacci and Robinson-Rechavi 2017; Yanai et al. 2005) was calculated for each gene/transcript in all seven vertebrates to determine the chamber specificity of each gene/transcript.

### Conserved enrichment networks

We produced combined enrichment plots for all the species with 4-chambered hearts. The enrichment networks were generated based on the DEGs in individual heart chamber comparisons. Enrichment networks were also produced based on the contrast of each individual chamber compared to the average expression of the other 3 chambers. The enrichment plots were generated using GSEA (v3.0) (Subramanian et al. 2005) based on the pre-ranked gene lists. All significantly enriched terms were then combined into functionally interpretable clusters using Enrichment Map (v3.2.1) (Merico et al. 2010) plugin in Cytoscape (v3.7.1) (Shannon et al. 2003).

### Identification of lncRNA homologs

The lncRNA homologs were identified based on sequence, secondary structure and syntenic position of the transcripts across the seven vertebrate species. For the sequence based lncRNA homologs identification, we employed an all-vs-all blastn based RBH search strategy based on the sequence of the lncRNAs and their promoter regions. OrthoMCL (v2.0.9) (Li et al. 2003), was then used to cluster these RBH hits into orthologous gene groups. To identify structure based lncRNA homologs, the secondary structures of lncRNAs were first determined using RNAFold program form ViennaRNA Package (v2.4.13) (Lorenz et al. 2011). We used Beagle (Mattei et al. 2015, 2014) to perform an all-vs-all alignment for all the species to identify the secondary structure based lncRNA homologs.

The synteny based lncRNA homologs were identified using the homologous neighboring PC genes. The syntenic homologs were classified into syntenic lncRNAs, if at least six of the ten PC neighbors were homologous. We also identified immediate syntenic lncRNA pairs based on immediate PC neighbors and classified both of these categories using strand information of the PC homologs.

### Determining the factors influencing the co-expression of neighboring transcripts

The co-expression values, for all the neighboring transcript pairs across human and mouse samples were calculated using the Spearman correlation coefficient. We contrasted the neighboring transcripts based on the overlap of the neighboring transcripts. We also looked at the impact of strandedness and the transcript biotype on the correlation value both for overlapping and non-overlapping transcripts. Finally, we also compared the correlation coefficient for conserved syntenic lncRNA-PC homolog pairs with non-conserved ones. All appropriate statistical analysis was performed in R (v3.5).

### Cardiac expressed circRNAs

The filtered RNA-Seq reads were mapped onto the reference genome using BWA-MEM (v0.7.17) (Li 2013) followed by circRNA detection using CIRI2 (v2.0.6) (Gao et al. 2018). Sailfish-cir (v0.11a) was used for the quantification of circRNA transcripts (Li et al. 2017). The differentially expressed circRNAs across all possible heart chambers were determined using DESeq2. CircTest was used to identify circRNAs whose expression is independent of their host gene expression values (Cheng et al. 2015). The conserved circRNAs were identified using the all-vs-all RBH strategy. The enrichment analysis was performed for the circRNA host genes using the online tool g:Profiler (Raudvere et al. 2019). The results for all organisms were then combined using the Enrichment Map plugin in Cytoscape.

### Data access

All raw and processed sequencing data generated in this study have been submitted to the National Center for Biotechnology Information (NCBI) under accession number PRJNA690979. The analysis results can be explored using the website portal http://evoactg.uni-muenster.de/.

## Acknowledgments

We would like to thank Eugenio Mattei for providing us with the local version of the Beagle software. We also thank the core facility genomics of the medical faculty at Münster, particularly the superb technical assistance by Elvira Barg and Marianne Jansen-Rust. The authors are grateful to Prof. Wolf-Michael Weber and Prof. Stefan Schulte-Merker for providing us with African frog and zebrafish heart biopsies respectively. The authors also acknowledge Dr. Jörn Scharsack and Yvonne Padberg for their help with the dissection of zebrafish heart. M.S. and S.G. are funded by the Deutsche Forschungsgemeinschaft (DFG, German Research Foundation) – RTG2220 – project number 281125614. U.S. and M.S. are funded by the Netherlands Heart Foundation (CVON2014-09). L.dW and M.S. are funded by the European Union’s Horizon 2020 research and innovation program under the Marie Sklodowska-Curie grant agreement no. 813716 (TRAIN-HEART Innovative Training Network).

