## Supplemental Materials for "Evolutionarily conserved transcriptional landscape of the heart defining the chamber specific physiology"

(1) Institute of Human Genetics, Division of Genetic Epidemiology, University of Muenster, Muenster, Germany. (2) Department of Physiology, Cardiovascular Research Institute Maastricht (CARIM), Maastricht University, Maastricht, The Netherlands (3) Department of Molecular Genetics, Maastricht University, Maastricht, The Netherlands (4) Department of Biochemistry, Genetic Epidemiology and Statistic al Genetics, Cardiovascular Research Institute Maastricht (CARIM), Maastricht University, Maastricht, The Netherlands (5) Department of Cardiothoracic Surgery, Cardiovascular Research Institute Maastricht (CARIM), Maastricht University Medical Centre+, Maastricht, The Netherlands.

**Contents:**

**Supplemental Methods	Page 2-7**

**Supplemental References	Page 8-9**

**Supplemental Methods**

**Tissue acquisition and processing**

All the study protocols dealing with the patients and tissue collection was performed in accordance to the ethical guidelines of the Helsinki Declaration. All necessary permits to use the listed samples for biomedical research were obtained and can be provided upon request.

The Casper strain of zebrafish heart chamber biopsies were obtained in triplicates from the Institute for Cardiovascular Organogenesis and Regeneration, WWU Münster (Prof. Stefan Schulte-Merker) where they were maintained according to FELASA recommendations (Aleström et al. 2020). Dissection of fish was performed at the same institute using the Olympus SZX16 microscope. The three African clawed frog heart samples were obtained from the Institute of Animal Physiology, WWU Münster (Prof. Wolf-Michael Weber). Both the animal tissues were obtained according to the guidelines of the animal ethics committees at the University of Münster, Germany. The two chicken and three B6129F1 mice biopsies were obtained from the CARIM School for Cardiovascular Diseases, Maastricht University (Prof. L.J. De Windt). The two New Zealand white rabbit strain and three Dutch white goat strain heart chamber biopsies were obtained from the CARIM School for Cardiovascular Diseases, Maastricht University (Prof. U. Schotten). The three human heart chamber biopsies were obtained as a part of the RACE V consortium from the same facility. Informed consent was obtained from all the human participants as a part of the RACE V consortium study. All the necessary approvals were taken from the ethical board for animal experimentation of Maastricht University were taken.

The samples were preferably collected from female only individuals to minimize the differences which might exist due to the gender of the organisms. We obtained all female heart chamber sample biopsies for mouse, goat, zebrafish and frog in triplicates. While for human samples, we obtained 2 male and 1 female samples. Additionally, for human samples, due to limitations in obtaining right ventricular biopsies due to ethical reasons, we only had samples for other 3 heart chambers. In the case of chicken and rabbit, the samples were collected in duplicates.

**Library preparation and transcriptome sequencing**

Total RNA was isolated using the Direct-zol RNA Microprep Kit (Zymo Research) including a DNase digestion step. The integrity of the total RNA was assessed by the means of Bioanalyzer RNA 6000 Nano Kit (Agilent) and then the RNA was used for rRNA depletion and subsequent directional library preparation (NEBNext Ultra II RNA Library Prep Kit, NEB). The quality of the resulting NGS library was determined by means of the Bioanalyzer High Sensitivity DNA Kit (Agilent). Equimolar library pools based on the library quantification results of the NEBNext Library Quant Kit for Illumina (NEB) were sequenced in a paired end mode 75 cycles on a NextSeq 500 system (Illumina) using v2 and v2.5 chemistry.

**Read mapping and transcriptome assembly**

The sequencing reads from each RNA-Seq library were subjected to stringent quality control. Using Trimmomatic (v0.36), adapters and all the reads with average Phred quality score < 20 were removed (Bolger et al. 2014). Poor quality bases were trimmed using the parameter SLIDINGWINDOW:4:15 and only reads longer than 35 bases were retained for further analysis. Processed reads from each sample were then aligned to respective species reference genome (Supplemental Table S1) using the STAR aligner (v2.5.3a) in 2-pass mapping mode which allows for unbiased exon splice junction detection (Dobin et al. 2013). In addition, the following parameters were modified while performing the 2nd pass alignment run: --outFilterType BySJout --alignSJoverhangMin 15 --alignSJDBoverhangMin 1 --outFilterMismatchNmax 33 --seedSearchStartLmax 12 --outFilterMatchNminOverLread 0 --outFilterScoreMinOverLread 0.3 --outSAMunmapped Within --outSAMattrIHstart 0 --outFilterIntronMotifs RemoveNoncanonicalUnannotated --outSAMtype BAM SortedByCoordinate. Mapped reads from each sample were independently assembled into transcripts using StringTie (v1.3.4d) guided by the reference genome annotations (Pertea et al. 2015). For each species, the annotations from individual samples were then merged into a single consensus genome annotation using the StringTie --merge option followed by the quantification of transcript abundance. One of the goat right atrial biopsies was not considered for further analysis due to low mapping percentage.

**Identification of novel cardiac expressed lncRNAs**

Novel transcript annotations for each species’ consensus genome were isolated and further analyzed to remove unreliable and spurious annotations. Firstly, all novel transcripts with length < 200 bases and FPKM > 0.5 in at least 2 samples were discarded, as these were most likely sequencing or assembly artifacts. All single exonic transcripts were then filtered out, due to the uncertainty associated with these assembled fragments. Next, CPC2 (v0.1) (Kang et al. 2017), CNCI (v2) (Sun et al. 2013), NCBI’s ORFfinder (v0.4.3) and Pfam database (RD et al. 2016) were employed to screen the candidate lncRNAs for evidence of coding potential. All the transcripts with CPC2 coding probability > 0.4, CNCI score >= 0 and having an ORF predicted by ORFfinder were filtered out. The remaining transcripts were translated in all three possible frames and PfamScan was used to identify any known protein family domains in Pfam database (release 31). Finally, any transcript with Pfam hit and E-value < 0.001 was discarded. Only high-confidence novel lncRNAs that were classified as non-coding by all the four tools were retained for further analysis.

**Expression quantification and differential expression analysis**

For each species, we merged our lncRNA annotations with the reference genome annotations (Supplemental Table S1). StringTie was used to quantify the expression of individual genes and transcript isoforms guided by the merged annotation file. The resulting read quantifications were then imported into R and DESeq2 (v1.22.2) (Love et al. 2014) was used to identify DEGs and DETs for all possible chamber comparisons within each species. Additionally, we checked for differential expression between atria and ventricles by combining the left and right halves in whichever species it was applicable. To identify DEGs and DETs, we independently performed differential expression analysis for each tissue comparison, both at the gene and transcript level. Genes and transcripts with total counts across samples less than 10 were excluded from the analysis. Only genes/isoforms with log2 fold change (FC) ≥ 1 or ≤ -1 and adjusted p-value < 0.05 (Benjamini-Hochberg) were considered as differentially expressed. Genes and transcripts with mean FPKM values ≥ 1 were defined as being expressed in the heart of that species. PCA plots were produced for each species using VST normalized reads in R.

**Chamber specificity**

To determine the heart chamber-specific expression, the τ index for each individual gene and transcript across all seven organisms was calculated (Kryuchkova-Mostacci and Robinson-Rechavi 2017; Yanai et al. 2005). The index was calculated only for those genes and transcripts, for which the total number of reads mapping exceeded 15 and 30 respectively. The value of the index varies between 0 and 1, with 1 being highly chamber specific and 0 being ubiquitously expressed. In the case of human samples, the index was calculated using the 3 available heart chambers. Similarly, the τ index was also calculated for circRNA transcripts with at least 15 reads mapping to them.

**Enrichment network analysis**

For each individual species, two different approaches were employed to create enrichment networks. In the first approach, DEGs in individual pairs of heart chamber were considered as the basis to create ranked gene lists. Through this approach, we were able to contrast the gene set modules operational across 2 chambers at a time. While, the second approach involved contrasting each individual chamber against the average expression of all the other available chambers for that species. For this, we used DESeq2 without an intercept and averaged out the expression of all other chambers while building contrasts. This approach, resulted in a list of gene set modules which are specific to that chamber as opposed to all the other chambers within that species. In both the strategies, genes were ranked based on the z-scores, which were calculated using adjusted *p*-values and signs of log fold change from DESeq2 results. GSEA (v3.0) (Subramanian et al. 2005) on the pre-ranked gene lists was performed using the gene set GMT (Gene Matrix Transposed) files derived from g:Profiler (Raudvere et al. 2019) using default parameters. For all species, except goats, the GMT files were downloaded based on Ensembl v90 gene IDs. In the case of goat, Ensembl v92 was used. Since no files were available for *X.laevis*, the GMT files for *X.tropicalis* were used. The Ensembl IDs in the GMT files were manually transformed to include *X.laevis* gene names based on Xenbase database annotations (Karimi et al. 2018). Next, we assembled all the significantly enriched terms into functionally interpretable clusters using Enrichment Map (v3.2.1) (Merico et al. 2010) plugin in Cytoscape (v3.7.1) (Shannon et al. 2003).

**Positional classification of lncRNAs**

For each species, the lncRNA coordinates from the merged annotation file were extracted and classified based on the genomic location with respect to the PC genes. For Ensembl annotations, all transcripts with following biotype were considered as lncRNAs: lincRNA, 3prime overlapping ncRNA, non-coding, antisense RNA, bidirectional promoter lncRNA, macro lncRNA, processed transcript, retained intron, sense intronic, and sense overlapping. Next, FEELnc (v1.0) classifier was applied with default parameters to segregate the lncRNAs based on the immediate PC transcripts within 100 kb (Kilobase) radius (Wucher et al. 2017). For each lncRNA, precedence was given to the immediate gene in case of intergenic assignment and to exonic over intronic annotation for overlapping annotations. LncRNAs with no coding transcripts located within 100 kb boundary were classified as isolated intergenic lncRNAs.

**Sequence based lncRNA homologs identification**

To discover sequence based lncRNA homologs, only intergenic and antisense lncRNA transcripts were considered. For each species, the sequence of all lncRNAs in FASTA format were extracted using gffread (v0.9.12). Next, an all-vs-all blastn search was performed to detect sequence similarity between lncRNAs across species. Only blast hits with identity >= 20% and E-value <= 10^-5^ were kept for further analysis. RBH strategy was applied to identify lncRNA pairs with best hits in two different species or best matching hits within the same organism. Finally, OrthoMCL (v2.0.9) (Li et al. 2003), which uses Markov clustering algorithm was employed to cluster these RBH hits into orthologous gene clusters. The same approach was also applied to identify lncRNA homologs based on the sequence similarity within their promoter region. The promoter region was defined as the region 2,000 bases upstream of lncRNA sequences. All lncRNAs with promoters shorter than 100 bases were removed. Using this approach, we were able to identify promoter based orthologous lncRNA families.

**Secondary structure based lncRNA homologs identification**

To determine structure based homologs, only lncRNAs shorter than 1,000 bases were considered. Using RNAFold program form ViennaRNA Package (v2.4.13) (Lorenz et al. 2011), the secondary structures for all the lncRNAs were computed across all seven species. Next, we used Beagle, which performs RNA secondary structure alignment using BEAR notations (Mattei et al. 2015, 2014). An all-vs-all alignment was performed for all the species and the results with Zscore significance of > 3 and structure identity ≥ 50 were filtered out.

**Synteny based lncRNA homologs identification**

In order to identify syntenic lncRNAs, homologous PC gene neighbors of the lncRNAs were used. Using the biomaRt package (Durinck et al. 2005) in R, the Ensembl ids (version 90) for all the homologous PC genes in chicken, humans, mouse, rabbit, and zebrafish were retrieved. For frog, the Ensembl ids (version 90) for *X.tropicalis* were retrieved, which were then mapped to *X.laevis* using the Xenbase database annotations (Karimi et al. 2018). Since the goat annotations were only made available from Ensembl version 92 onwards, the homologous gene ids for other species were retrieved for version 92 and manually converted to version 90. Based on these homologous genes, we organized them in homologous groups, wherein each group represents all possible homologous PC genes across all the seven species.

Next, for each lncRNA, five upstream and downstream homologous PC genes were considered to detect positional conservation. For simplicity, the overlapping PC genes were also labelled as up/downstream based on their start site. LncRNAs with less than 3 neighboring genes either side, were not considered for the analysis. Based on the homologous groups, group ids were assigned to the neighboring genes and syntenic homologs were found using 2 strategies: (i) lncRNAs with at least 6 out of 10 neighboring coding genes with same group ids were considered as syntenic homologs; (ii) lncRNAs with immediate neighboring genes having the same group ids were called immediate syntenic homologs. These lncRNA homologs were further classified on the basis of the strand orientation of neighboring genes into stranded syntenic homologs and immediate stranded syntenic homologs.

**Co-expression of neighboring gene pairs**

To look at the co-expression values, all the lncRNA and PC transcripts across human and mouse samples were considered. For both the species, an all-vs-all FEELnc (v1.0) classifier run was performed, to look for all possible overlaps for these transcripts. Using this approach, we were able to determine all possible overlaps for lncRNA and PC transcripts. All those transcripts with no overlap in the 100 kb vicinity were removed from further analysis Based on the amount of overlap between transcripts, the resulting transcript pairs were then classified into non-overlapping neighboring transcripts or overlapping transcript pairs. Next, to look at the effect of strandedness of the transcript pairs, were further classified the transcript pairs based on if both the transcripts were present on the same or opposite strand. Finally, the transcript pairs were divided based on the biotypes of transcripts involved (lncRNA/PC). The read count data for transcripts in each species was pre-filtered using total reads > 10 and then transformed using VST in DESeq2. Spearman correlation coefficient was calculated individually for all transcript pairs in each category. Wilcoxon signed-rank test was then performed to test for significant differences between distinct categories for human and mouse samples. For the biotype based comparison, one-way Kruskal-Wallis test was undertaken followed by Dunn’s test to analyze the multiple group comparisons for all the three possible transcript pair biotypes.

**Identification and quantification of cardiac circRNAs**

For each species, the filtered reads obtained after running Trimmomatic, were mapped onto their respective reference genomes (Supplemental Table S1) using BWA-MEM (v0.7.17) with the parameter -T= 19 (Li 2013). CIRI2 (v2.0.6) was used for the genome-wide detection and annotation of all putative circRNAs (Gao et al. 2018). It utilises the paired chiastic clipping signals to help identify junctional reads which form the circRNAs. Only circRNAs which are expressed in 50% of the samples in each species and have back-splicing read sum > 10 mapping to them were considered for further analysis.

Sailfish-cir (v0.11a) was used for the quantification of circRNA transcripts (Li et al. 2017). Sailfish-cir transforms the circular transcripts into pseudo-linear transcripts followed by model-based quantification using Sailfish (Patro et al. 2014). For each species, the circRNA counts were then used to identify differentially expressed circRNAs across all possible heart chambers using DESeq2. CircRNAs with log2 fold change (FC) >= 1 or <= -1 and adjusted p value < 0.05 (Benjamini-Hochberg) were considered differentially expressed. CircTest package was used to identify circRNAs whose expression is independent of their host gene expression values (Cheng et al. 2015).

**Conservation of circular RNAs**

To identify conserved circRNAs, the sequence conservation across all the seven species was used. For exonic circRNAs, only the FASTA sequence of the exons were extracted, while for intronic and intergenic circRNAs, the whole sequence was considered. Next, an all-vs-all RBH strategy was applied to discover circRNA homologs. The blast search parameters were the same as applied for lncRNA homologs identification.

**Enrichment network of circRNA host genes**

To explore the processes in which circRNAs are involved, enrichment analysis was performed for each species using all the genes coding for circRNAs transcripts. Next, pathway analysis was performed using the online tool g:Profiler (Raudvere et al. 2019) with default parameters and pathways with term size between 10 and 1000 were selected for further analysis. Finally, the enrichment results for all organisms were combined and imported into Cytoscape using the Enrichment Map plugin.

**Website portal**

A web resource, called EvoACTG - Evolutionary Atlas of Cardiac Transcriptome and homologous gene was also developed. The database provides the users with information about the expression of homologous genes in various heart chambers across all the seven species. The chamber specificity module was developed using shiny package in R. We also developed a homolog browser in shiny using the TnT genome browser (Pignatelli 2016). The website is available via the link (<http://evoactg.uni-muenster.de/>).
