## Supplemental Figure Legends for "Evolutionarily conserved transcriptional landscape of the heart defining the chamber specific physiology"

**Supplemental Fig. S1: Distribution of lncRNA annotation across the seven vertebrate species.**

The graph shows the number and the percentage of annotated and novel lncRNAs identified in the heart chambers of zebrafish, frog, chicken, goat, human, mouse and rabbits.

**Supplemental Fig. S2: Expression profiles across various heart chambers.**

Principal Component analysis of VST normalised read counts for top 500 cardiac expressed genes for all the seven vertebrates including human dataset from Johnson et. al.

**Supplemental Fig. S3: Several genes express in a chamber specific manner.**

(A) The distribution of the number of chamber specific genes across all the seven vertebrates. (B) The distribution of the biotypes of chamber specific genes in all seven species.

**Supplemental Fig. S4: Enrichment modules enriched for heart chamber comparisons.**

Enrichment terms significantly (P < 0.05) enriched in (A) LA vs RA (B) LV vs RV (C) LA vs LV and (D) RA vs RV were generated using the Enrichment map for all vertebrates with four chambered hearts. The human dataset from Johnson et al. was also included in the analysis. Each node represents individual enriched term with the parts of the pie chart representing enrichment in individual organisms. The degree of enrichment is represented by the colour for each individual comparison. Clusters of similarly enriched terms are indicated by circles with consensus term labels.

**Supplemental Fig. S5: Enrichment modules enriched in individual heart chambers.**

Enrichment terms significantly (P < 0.05) enriched in (A) LA (B) LV (C) RA and (D) RV compared to the average expression across all other chambers were generated using the Enrichment map for all vertebrates with four chambered hearts. The human dataset from Johnson et al. was also included in the analysis. Each node represents individual enriched term with the parts of the pie chart representing enrichment in individual organisms. The degree of enrichment is represented by the colour for each individual comparison. Clusters of similarly enriched terms are indicated by circles with consensus term labels.

**Supplemental Fig. S6: Sequence conservation of syntenic lncRNAs conserved between human and mouse.**

The volcano plot compares the percentage of sequence identity for the various sub-categories of conserved human-mouse syntenic lncRNA pairs. There is significant difference (Wilcox test) between the percentage identity for immediate stranded (p-value=1.45e-31), immediate unstranded (p-value=2.52e-32), stranded (p-value=4.37e-58) and unstranded (p-value=5.87e-41) conserved syntenic human-mouse lncRNA pairs as compared to random human-mouse non-syntenic lncRNAs.

**Supplemental Fig. S7: Expression modules enriched for circRNA host genes across the hearts of seven vertebrates.**

The diagram shows the various enrichment terms significantly (P < 0.05) enriched based on the genes that produce circRNAs within the heart. Each node represents individual enriched term with the parts of the pie chart representing enrichment in individual organisms. Clusters of similarly enriched terms are indicated by circles with consensus term labels.
