## Supplementary figures and images for "Evolutionarily conserved transcriptional landscape of the heart defining the chamber specific physiology"

### Supplemental Figure S1

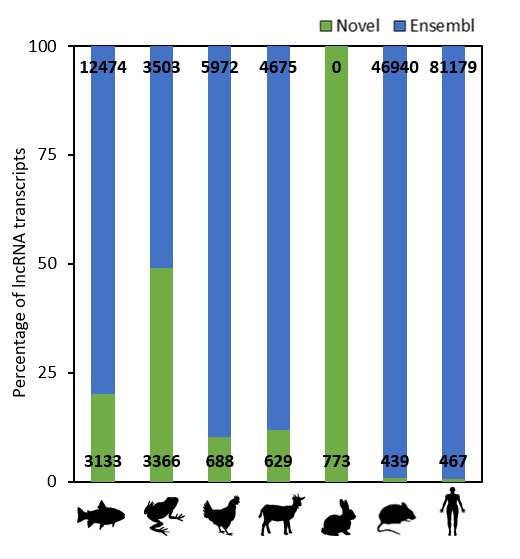

### Supplemental Figure S2

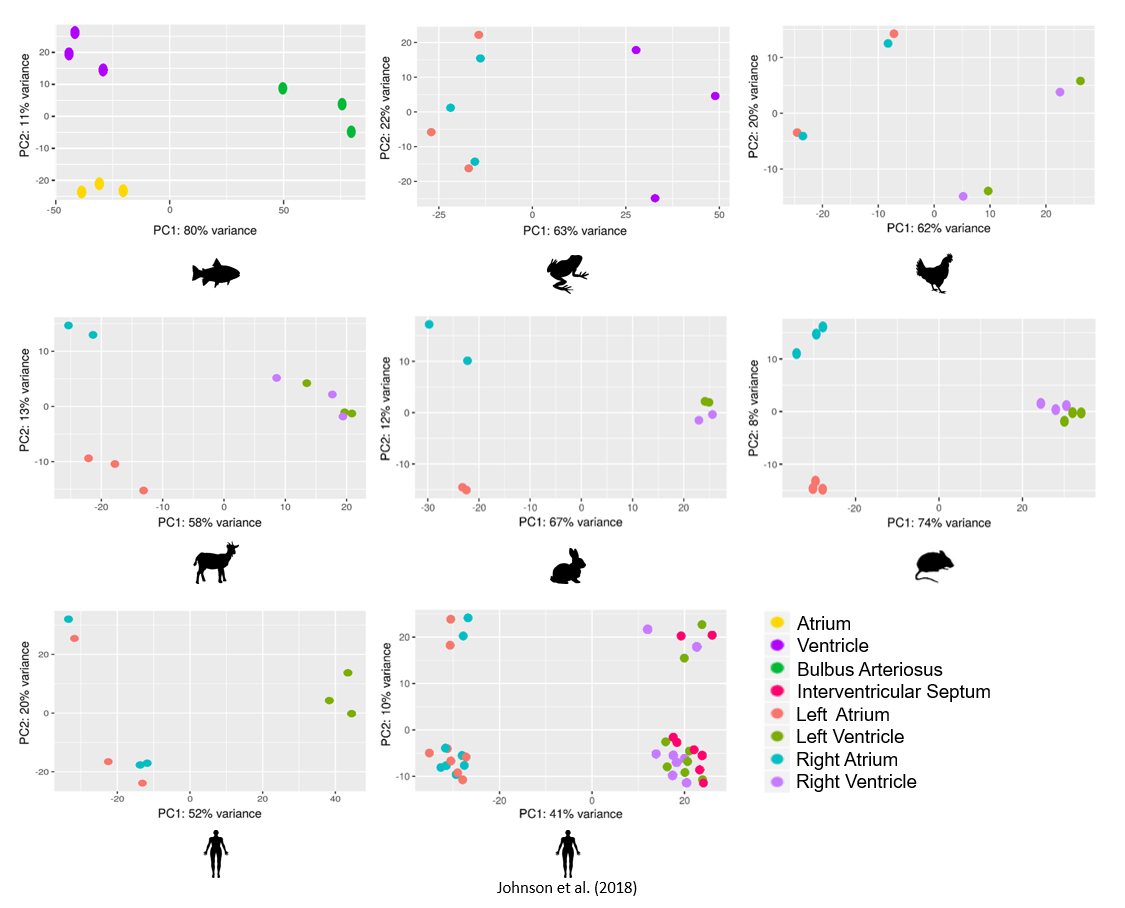

### Supplemental Figure S3

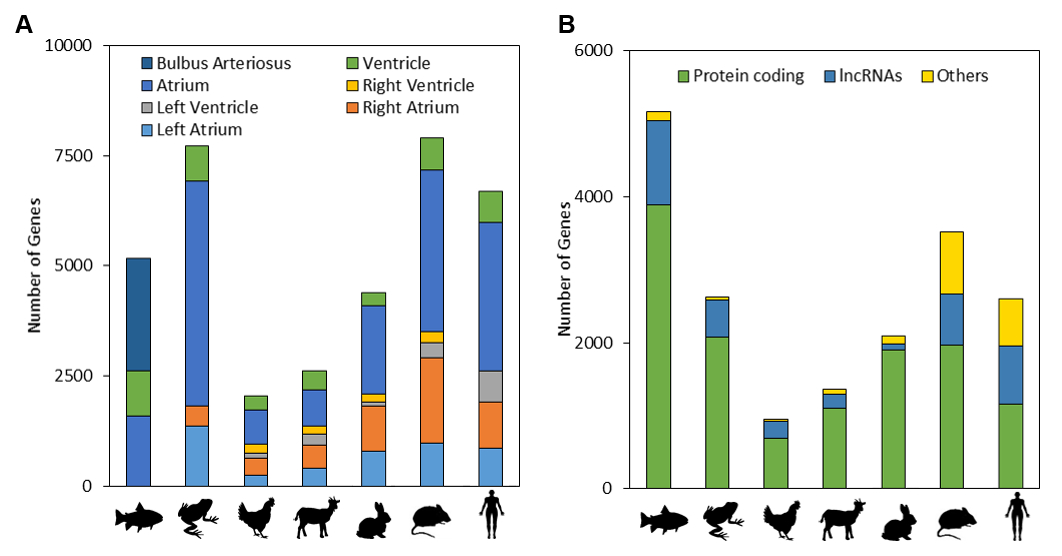

### Supplemental Figure S4

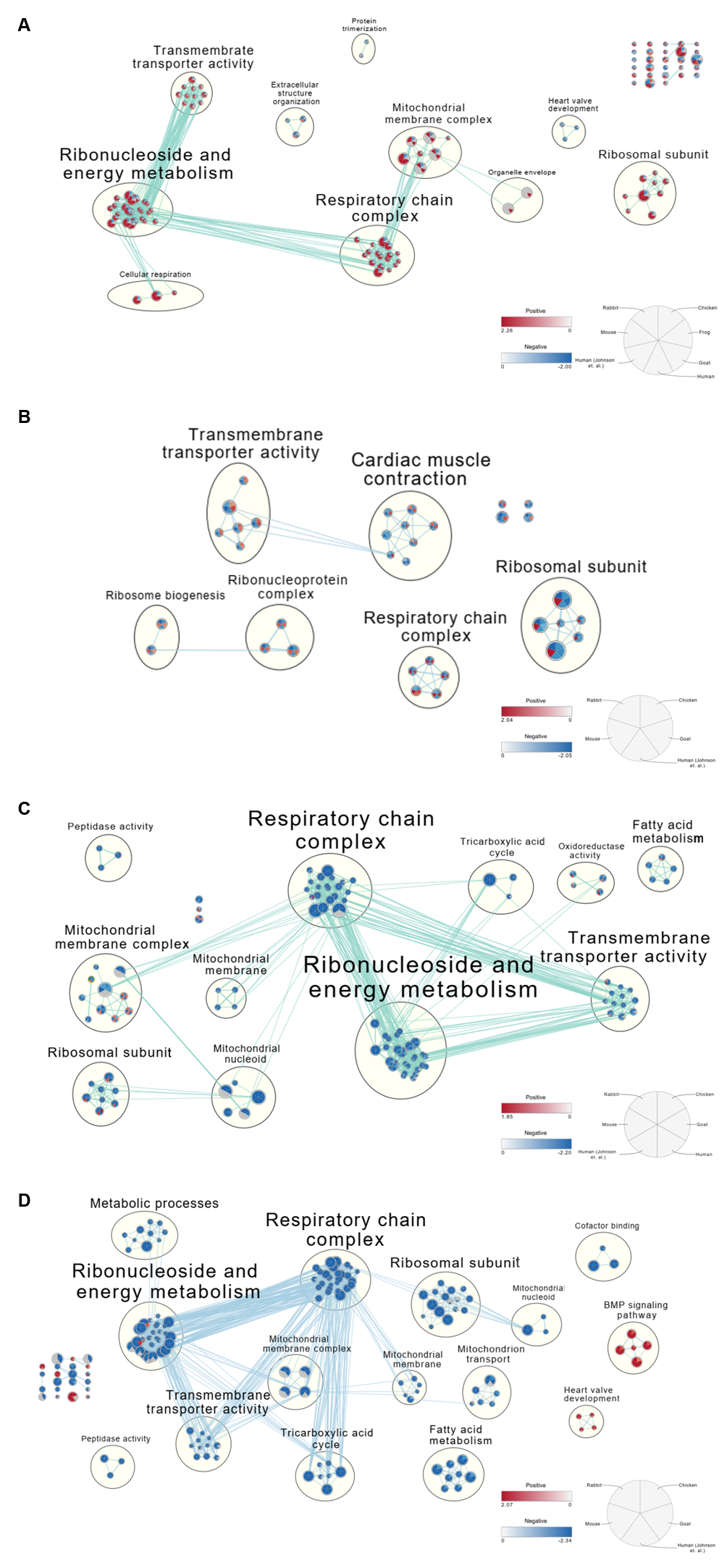

### Supplemental Figure S5

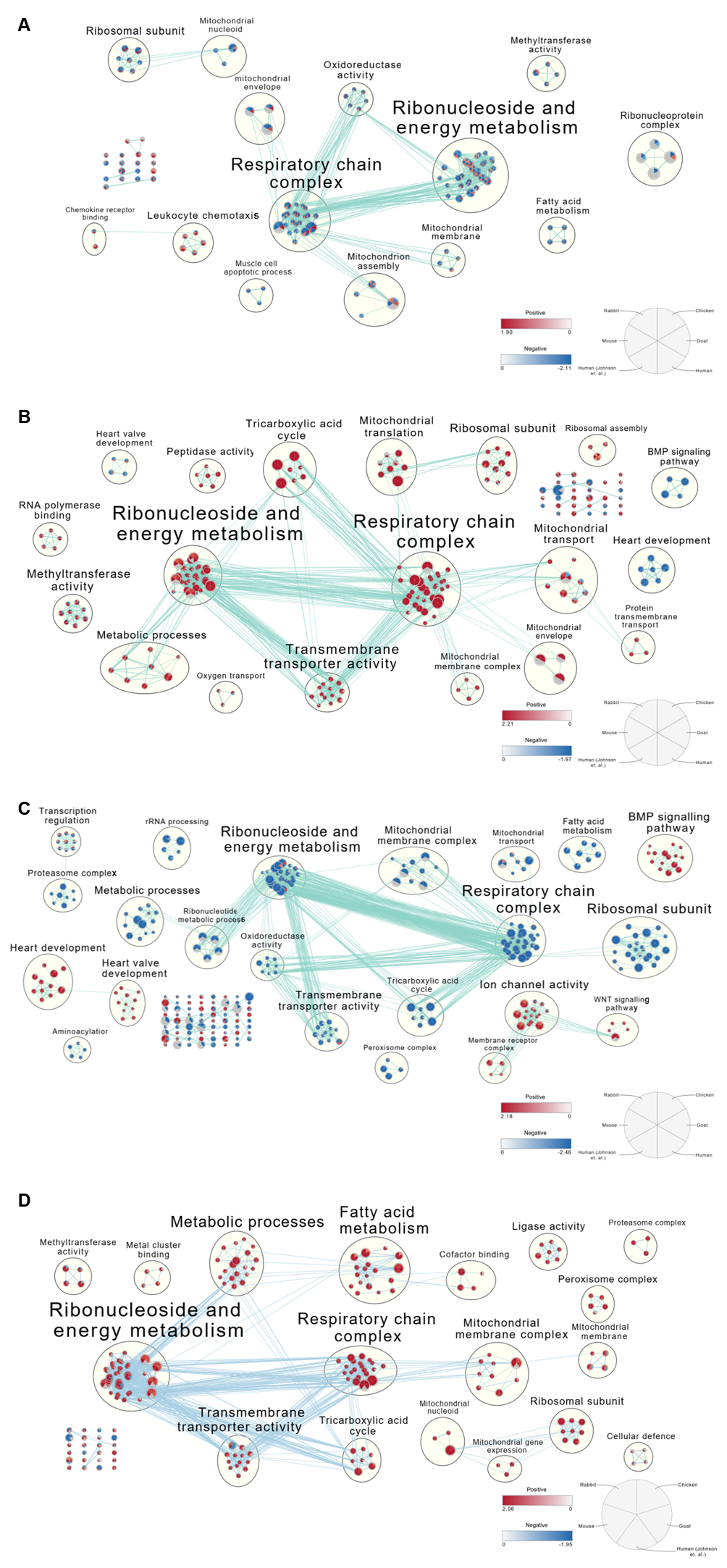

### Supplemental Figure S6

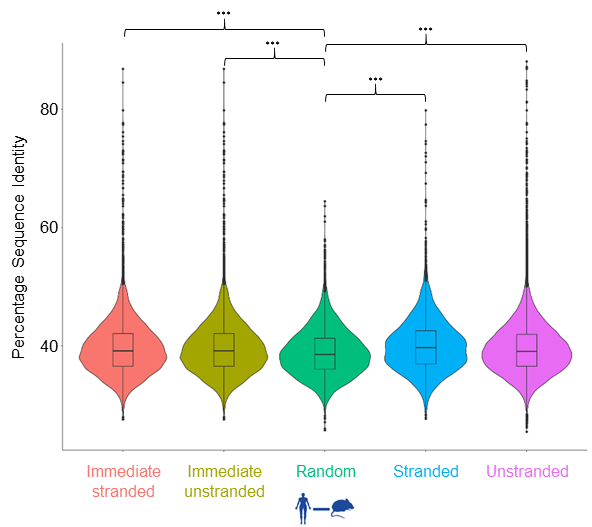

### Supplemental Figure S7

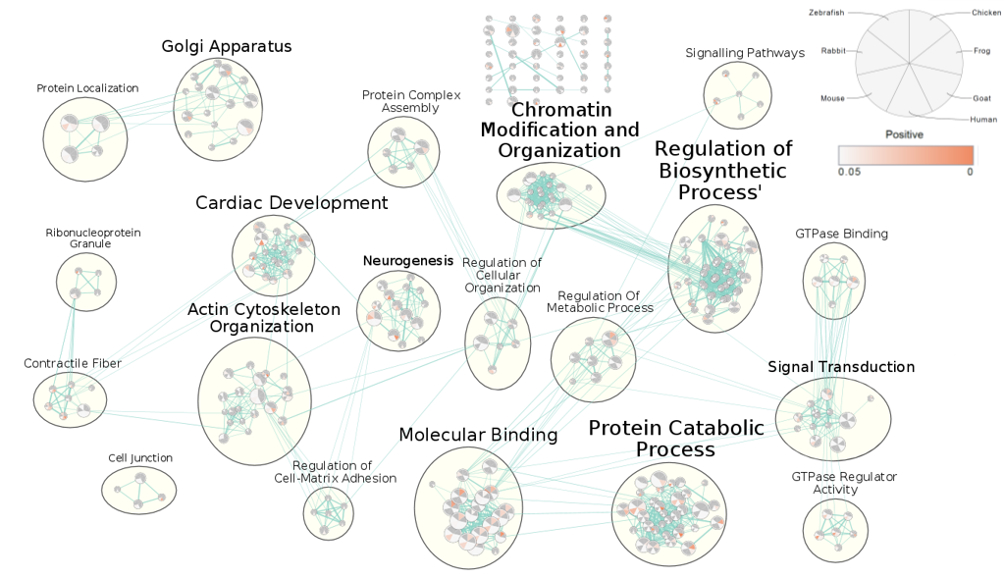
